# *Aedes aegypti* (Aag2)-derived clonal mosquito cell lines reveal the impact of pre-existing persistent infection with the insect-specific bunyavirus Phasi Charoen-like virus on arbovirus replication

**DOI:** 10.1101/596205

**Authors:** Anthony C. Fredericks, Louisa E. Wallace, Tiffany A. Russell, Andrew D. Davidson, Ana Fernandez-Sesma, Kevin Maringer

**Affiliations:** Department of Microbiology, Icahn School of Medicine at Mount Sinai, New York, NY USA; Department of Microbial Sciences, Faculty of Health and Medical Sciences, University of Surrey, Guildford, United Kingdom; School of Cellular and Molecular Medicine, University of Bristol, Bristol, United Kingdom

## Abstract

**Background:** *Aedes aegypti* is a vector mosquito of major public health importance, transmitting arthropod-borne viruses (arboviruses) such as chikungunya, dengue, yellow fever and Zika viruses. Wild mosquito populations are persistently infected at high prevalence with insect-specific viruses that do not replicate in vertebrate hosts. In experimental settings, acute infections with insect-specific viruses have been shown to modulate arbovirus infection and transmission in *Ae. aegypti* and other vector mosquitoes. However, the impact of persistent insect-specific virus infections that more closely mimic the situation in nature has not been investigated extensively. Cell lines are useful models for studying virus-host interactions, however the available *Ae. aegypti* cell lines are poorly defined and heterogenous cultures.

**Methodology/Principle Findings:** We generated single cell-derived clonal cell lines from the commonly used *Ae. aegypti* cell line Aag2. Two of the fourteen Aag2-derived clonal cell lines generated harboured markedly and consistently reduced levels of the insect-specific bunyavirus Phasi Charoen-like virus (PCLV) known to persistently infect Aag2 cells. In contrast to studies with acute insect-specific virus infections in cell culture and *in vivo*, we found that pre-existing persistent PCLV infection had no major impact on the replication of the flaviviruses dengue virus and Zika virus, the alphavirus Sindbis virus, or the rhabdovirus vesicular stomatitis virus. We also performed a detailed characterisation of the morphology, transfection efficiency and immune status of our Aag2-derived clonal cell lines, and have made a clone that we term Aag2-AF5 available to the research community as a well-defined cell culture model for arbovirus-vector interaction studies.

**Conclusions/Significance:** Our findings highlight the need for further *in vivo* studies that more closely recapitulate natural arbovirus transmission settings in which arboviruses encounter mosquitoes harbouring persistent rather than acute insect-specific virus infections. Furthermore, we provide the well-characterised Aag2-derived clonal cell line as a valuable resource to the arbovirus research community.

**AUTHOR SUMMARY:** Mosquito-borne viruses usually only infect humans through the bite of a mosquito that carries the virus. Viruses transmitted by the ‘yellow fever mosquito’ *Aedes aegypti*, including dengue virus, Zika virus, yellow fever virus and chikungunya virus, are causing an ever-increasing number of human disease cases globally. Mosquito-borne viruses have to infect and replicate inside the mosquito before they are transmitted to humans, and the presence of other infectious agents can change the efficiency of virus transmission. Mosquitoes are known to be infected with ‘insect-specific viruses’ that only infect mosquitoes and cannot cause human disease. We have shown here that in laboratory cell cultures derived from the *Aedes aegypti* mosquito, pre-existing infection with an insect-specific virus called Phasi Charoen-like virus does not affect the infection and growth of the mosquito-borne viruses dengue virus, Zika virus, Sindbis virus or vesicular stomatitis virus. Compared to previous research, our research is more reflective of conditions that mosquito-borne viruses encounter in nature, and our results provide important new insights into whether and how insect-specific viruses affect mosquito-borne virus transmission. Ultimately, this information could inform ongoing research into whether insect-specific viruses could be used to prevent the transmission of mosquito-borne viruses to reduce global disease burdens.

## INTRODUCTION

Arthropod-borne viruses (arboviruses) are a major public health concern worldwide, with many considered emerging or re-emerging pathogens [1]. Significant taxons to which arboviruses belong include the positive-sense single-stranded RNA (+ssRNA) families *Flaviviridae* (genus *Flavivirus*) and *Togaviridae* (genus *Alphavirus*), and the negative-sense single-stranded RNA (-ssRNA) order *Bunyavirales* and family *Rhabdoviridae* (genus *Vesiculovirus*). Many arboviral taxons also include related insect-specific viruses that can infect vector insects but not vertebrate hosts [2,3]. Arboviruses transmitted by the vector mosquito *Aedes aegypti* are of particular concern to human health, as this mosquito species thrives in urban environments and is highly anthropophilic, feeding primarily on humans [4]. *Ae. aegypti* is the primary vector for the emerging and re-emerging flaviviruses dengue virus (DENV), yellow fever virus (YFV) and Zika virus (ZIKV), and the alphavirus chikungunya virus (CHIKV) [5].

Vector competence is the intrinsic ability of an arthropod to be infected with and transmit vector-borne pathogens [6]. Vector competence varies between individuals and populations based on many factors, including the combination of pathogen and vector genotype, co-infection status of the vector with other microbes, and other environmental factors [4,7-9]. There is widespread interest in understanding the underlying mechanisms influencing vector competence to gain a better understanding of how arboviruses are transmitted and emerge on a global and local scale, especially because this knowledge could aid the development of mosquitoes unable to transmit arboviruses of human public health concern. For example, mosquitoes harbouring the obligate intracellular bacteria *Wolbachia spp.* are less able to transmit DENV and other arboviruses [10-13] and are being released in endemic settings to test their impact on human disease burdens [14]. Similarly, insect-specific viruses have also been proposed as potential biocontrol agents to reduce arbovirus transmission [2,15].

Insect-specific viruses are highly prevalent in wild mosquito populations [16-28], with a number of studies investigating whether insect-specific viruses influence vector competence [reviewed in 15]. There is no consensus on how insect-specific viruses affect arbovirus replication in tissue culture or *in vivo*, with the experimental outcome varying depending on the combination of arbovirus, insect-specific virus and mosquito species (and potentially the specific mosquito line or cell line used), as well as other variations in the experimental set up [15]. Thus, previous studies have found insect-specific viruses to either increase [29,30], decrease [19,31-40] or have no effect [31,35,40-42] on the replication of various arboviruses across different mosquito species and cell lines. The majority of these studies were performed in the context of acute insect-specific virus infection, which may not accurately recapitulate the effects of the persistent insect-specific virus infections more commonly encountered in nature.

To our knowledge, there are no *in vivo* studies on the impact of persistent insect-specific virus infection on arbovirus replication in *Aedes spp.* Of the *in vitro* studies that tested arbovirus replication in *Aedes spp.*-derived cell lines in which persistent insect-specific virus infection was maintained over multiple cell passages, Burivong *et al.* found that DENV replication was reduced in the presence of persistent but not acute co-infection with Aedes albopictus densovirus (AalDV; family *Parvoviridae*) in the *Ae. albopictus* cell line C6/36 [40]. This highlights the differential effects persistent and acute insect-specific virus infections can have on arbovirus replication. Meanwhile, Fujita *et al.* showed that persistent infection with Shinobi tetravirus (SHTV; family *Permutotetraviridae*) and Menghai rhabdovirus (MERV; family *Rhabdoviridae*) alone and in combination reduce ZIKV replication in C6/36 cells, with the two viruses combined also suppressing the replication of the flaviviruses DENV and Japanese encephalitis virus (JEV) [39]. Both studies may not fully recapitulate the conditions encountered in nature, since C6/36 cells are immunocompromised [43], and the mosquito immune system is known to pose a barrier to arboviral infection, within-vector dissemination and transmission [44-48], and is a known contributor to the observed variability in vector competence [44,47]. Notably, Parry and Asgari observed only a modest reduction in DENV replication in the *Ae. aegypti* Aa20 cell line in the presence of Aedes anphevirus (AeAV; order *Mononegavirales*) [19]. Therefore, further studies into whether and how persistent infection with insect-specific viruses might modulate arbovirus replication *in vivo* or in immunocompetent *Aedes spp.* cell lines are needed.

The *Ae. aegypti*-derived cell line Aag2 is one of the most commonly used cell lines for studies into virus-vector interactions in tissue culture. One of the benefits of Aag2 cells is that they are immunocompetent [49,50]. A major antiviral immune response in mosquitoes is the RNA interference (RNAi) pathway, in which viral double-stranded RNAs (dsRNAs) are processed by Dicer-2 into small interfering RNAs (siRNAs) that are loaded into the RNA-induced silencing complex (RISC) to target and degrade viral RNAs and thus reduce viral replication and spread [51]. In addition, innate immune pathways such as the Janus kinase-signal transducer and activator of transcription (Jak-STAT) pathway and nuclear factor kappa-light-chain-enhancer of activated B cells (NF-κB)-regulated Toll and immunodeficiency (IMD) pathways regulate the expression of antimicrobial peptides that are induced upon microbial stimulation [51].

The Aag2 cell line was originally generated in the 1960’s by Peleg from whole homogenised embryos, and has been referred to as ‘Aag2’ since the 1990’s when Lan and Fallon adapted the culture for growth in E-5 medium [52]. Cells within the culture exhibit differing morphologies (Fig 1A), and it has been suggested that the varying morphologies of mosquito cells in culture may be indicative of the presence of a diversity of embryonic and differentiated cell types [53,54]. Furthermore, Aag2 cells are known to be persistently infected with a number of insect-specific viruses. Cell fusing agent virus (CFAV; family *Flaviviridae*, genus *Flavivirus*) was the first insect-specific virus discovered and has long been known to persistently infect Aag2 cells and other *Ae. aegypti* cell lines [3,55,56]. In addition, we previously discovered Aag2 cells to be persistently infected with the insect-specific virus Phasi Charoen-like virus (PCLV; order *Bunyavirales*, family *Phenuiviridae*, genus *Phasivirus*) [57]. CFAV and PCLV both circulate in *Ae. aegypti* in the wild [16,17,25], and may have entered the cell line during its establishment or later on from an infected laboratory mosquito colony or environmental sample. While some research groups have found their Aag2 cell lines to also be persistently infected with the insect-specific viruses AeAV [19] or Culex Y virus [58], this is not the case for our Aag2 cells [57].

**Fig 1.**
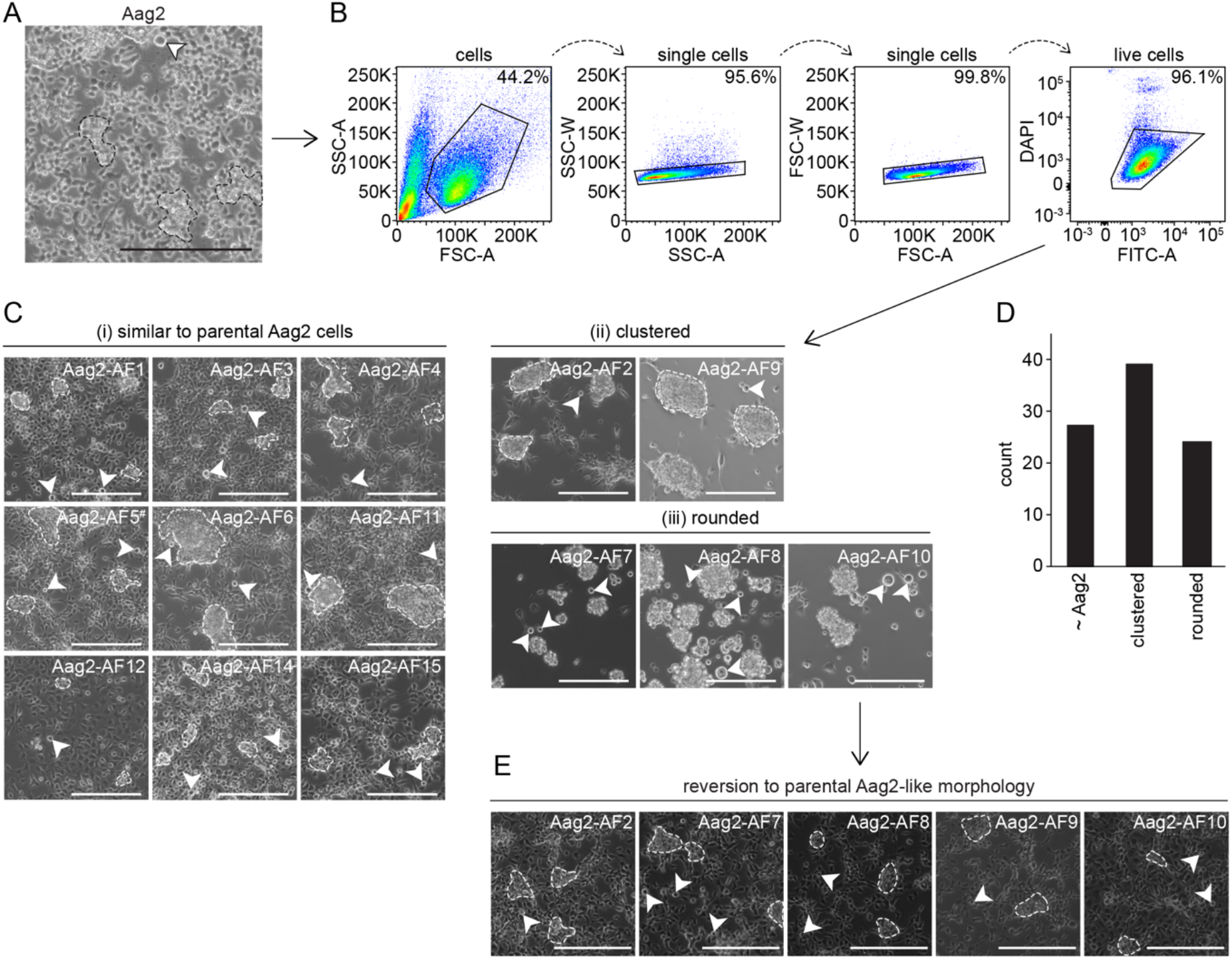
Generation of clonal Aag2-derived cell lines originating from single cells. (A) Brightfield microscopy image of heterogeneous Aag2 cell population consisting of multicellular ‘clusters’ (examples indicated by hashed lines throughout) and large rounded floating cells (arrows) interspersed across a loose monolayer. (B) FACS gating strategy illustrating selection of live single cells from DAPI-stained Aag2 cell suspension. (C) Resultant Aag2-derived clonal cell line morphologies following limited expansion; (i) similar appearance to parental Aag2 cells, (ii) highly clustered cells with no monolayer formation (some rounded floating cells present), (iii) only large rounded floating cells observable (individual cells and large multi-cell floating aggregates). Only those fourteen clones selected for further study are shown. Images were taken immediately following three-week expansion from single cells into confluent 24-well plate culture. ^#^ The Aag2-AF5 cell line was selected for CRISPR gene editing [59,60] (see main text). (D) Total number of clonal cell lines of each morphology generated. (E) Reversion of ‘clustered’ and ‘rounded’ clonal cell lines back to parental Aag2-like morphology following extended culture. Scale bar is 200 µm.

Here, we single-cell sorted Aag2 cells to generate clonal Aag2-derived cell lines so as to provide a better-defined homogeneous Aag2-derived cell line for the research community, and as a starting point for our own CRISPR experiments [59,60]. Although we initially selected clonal cell lines exhibiting different morphologies corresponding to those observed within the original Aag2 cell line (herein referred to as the ‘parental’ Aag2 cell line), these morphologies were not stable and all cell lines reverted to the parental Aag2 cell morphology. This suggests that the various cell morphologies observed in Aag2 cell cultures do not represent fundamentally different cell types. Furthermore, two of the clones selected for further characterisation were found to be ‘cured’ of PCLV, or at least harboured markedly and consistently reduced levels of PCLV. We used these clones to test the effect persistent (rather than acute) insect-specific virus infection has on superinfection with DENV, ZIKV, Sindbis virus (SINV) and vesicular stomatitis virus (VSV), and observed no notable reproducible impact on the replication of these arboviruses. Finally, we characterised a clone we termed ‘Aag2-AF5’ in further detail, and have provided this to the research community as a more well-defined version of the parental Aag2 cell line via the European Collection of Authenticated Cell Cultures (ECACC) (phe-culturecollections.org.uk). Our findings provide important insights into the impact that insect-specific viruses have on mosquito vector competence for arboviruses in an experimental set up that more accurately mimics conditions encountered by arboviruses in nature, with implications for the potential use of insect-specific viruses as biocontrol agents for reducing arbovirus transmission. Furthermore, our single cell-derived clone Aag2-AF5 represents a much needed standardised and well-defined *Ae. aegypti* cell line that will benefit the vector research community.

## METHODS

### Cells

Aag2 cells were a kind gift from Raul Andino (University of California, San Francisco, CA USA), and were maintained in Leibovitz’s L-15 medium supplemented with 2 mM glutamine (Sigma-Aldrich, St. Louis, MO USA), 0.1 mM non-essential amino acids (Sigma-Aldrich), 10% (v/v) tryptose phosphate broth (Sigma-Aldrich), 100 U/ml penicillin, 100 μg/ml streptomycin and 10% (v/v) foetal bovine serum (FBS) at 28°C in a humidified atmosphere without CO_2_. In our hands, the source of FBS is critically important for culturing Aag2 cells and derived clones, with ThermoFisher Scientific (Waltham, MA USA) product number 94000014 being optimal. C6/36 cells were a kind gift from Jorge Munoz-Jordan (Centers for Disease Control and Prevention, San Juan, Puerto Rico), and were maintained in Roswell Park Memorial Institute (RPMI) medium supplemented with 0.15% (w/v) sodium bicarbonate (Sigma-Aldrich), 0.1 mM non-essential amino acids, 2 mM L-glutamine, 1 mM sodium pyruvate and 10% (v/v) FBS at 33°C in a humidified atmosphere with 5% CO_2_. Baby hamster kidney (BHK) cells were a kind gift from Sujan Shresta (La Jolla Institute for Allergy and Immunology, La Jolla, CA USA), and were maintained in minimal essential medium (α-MEM) GlutaMAX supplemented with 10% (v/v) FBS, 100 U/ml penicillin, 100 μg/ml streptomycin, and 10 mM HEPES at 37°C in a humidified atmosphere with 5% CO_2_. Unless stated, reagents were from ThermoFisher Scientific. Madin Darby Canine Kidney (MDCK) cells were obtained from the American Type Culture Collection (ATCC) (Manassas, VA USA).

### Viruses

DENV serotype 2 (DENV-2) strain 16681 [61] was a kind gift from Richard Kinney (Arbovirus Disease Branch, Centers for Disease Control and Prevention, Fort Collins, CO USA). ZIKV strain MR766 [62] was obtained from ATCC. Green fluorescent protein (GFP)-expressing SINV, based on clone dsTE12Q [63,64], was a kind gift from Christopher Basler (Georgia State University, Atlanta, GA USA). VSV was the Indiana strain and expresses GFP [65], and was a kind gift from Adolfo Garcia-Sastre (Icahn School of Medicine at Mount Sinai, New York, NY USA). GFP-expressing Newcastle disease virus (NDV), based on clone Hitchner B1 [66], was a kind gift from Christopher Basler. DENV-2 and ZIKV were routinely grown on C6/36 cells at 33°C, and SINV and VSV were routinely grown on BHK cells at 37°C, in cell culture medium supplemented with 2% (v/v) FBS. Briefly, for ZIKV, SINV and VSV, confluent cell monolayers were infected at multiplicity of infection (MOI) 0.05 one day post-seeding. Culture supernatant was harvested seven (ZIKV) or two (SINV, VSV) days post-infection and clarified by centrifugation before storage, titration and use in experiments. For DENV-2, C6/36 cells were seeded at 1 × 10^6^ cells per 75 cm^2^ culture flask and infected one day later at MOI 0.5; virus was harvested seven days post-infection as described above.

For one-step growth curves, Aag2 cells were seeded at 5 × 10^5^ cells/well in 12-well plates (DENV-2), 3 × 10^5^ cells/well in 24-well plates (SINV, VSV) or 1 × 10^5^ cells/well in 96-well plates (ZIKV) and infected the next day by replacing culture medium with inoculum in phosphate-buffered saline (PBS). Inoculum was removed after 1 h and cells were washed once in PBS before adding fresh culture medium. To compare virus replication in Aag2-derived clonal cell lines, cells were seeded at 1 × 10^5^ cells/well in 96-well plates and infections performed as above. All viruses were titrated on confluent BHK cells one day post-seeding in culture medium containing 2% (v/v) FBS and 0.5% (w/v) methyl cellulose (Sigma Aldrich). For DENV-2 titrations, cells were moved to 33°C during and following infection. Titrations were fixed six (DENV-2) or three (ZIKV, SINV, VSV) days post-infection in 1% crystal violet solution in 20% ethanol following removal of the methyl cellulose overlay.

### Bacteria

*Escherichia coli* DH5α (ThermoFisher Scientific) were cultured overnight at 37°C with shaking in Luria Bertani (LB) medium (Sigma Aldrich) without antibiotics, and titrated on LB agar in 6-well plates overnight at 37°C. *Listeria monocytogenes* cultures were a kind gift from Adolfo Garcia-Sastre and *Staphylococcus aureus* cultures were a kind gift from Flora Samaroo (Icahn School of Medicine at Mount Sinai); both were cultured and titrated in brain heart infusion broth/agar (Sigma Aldrich) as for *E. coli*. Bacteria were pelleted, washed once in PBS and resuspended in PBS before heat-inactivation at 60°C for 3 h (*E. coli*, *L. monocytogenes*) or at 75°C for 6 h (*S. aureus*).

### Plasmids

The GFP expression vector pIEx-EGFP [67] was a kind gift from Doug Brackney (The Connecticut Agricultural Experiment Station, New Haven, CT USA). The constitutive firefly luciferase expression plasmid pKM19 was generated by amplifying the firefly luciferase gene from pLUC-MCS (Agilent Technologies, Santa Clara, CA USA) and cloning it into pIEx-EGFP after the enhanced GFP (EGFP) sequence was removed by digestion with XhoI and NcoI, using In-Fusion cloning (Takara Biosciences, Mountain View, CA USA). The constitutive *Renilla* luciferase expression plasmid pKM50 was generated by amplifying the *Ae. aegypti* ubiquitin UbL40 promoter from pSLfa-UbL40-EGFP [68] (a kind gift from Raul Andino) and cloned by In-Fusion into pRL-TK-Renilla (Promega, Madison, WI USA) after the TK promoter was removed by digestion with BglII and BstBI. pKM19 and pKM50 have been made available via Addgene (addgene.org, Watertown, MA USA) with reference numbers 123655 and 123656 respectively.

### Single Cell Sorting of Aag2 Cells

Prior to cell sorting, Aag2 cells were grown in Leibovitz’s L-15 medium with supplements as described above, including 20% (v/v) FBS. Adherent cells were trypsinised, pelleted by centrifugation, washed once in PBS, pelleted again by centrifugation and resuspended at 3−5 × 10^6^ cells/ml in sterile PBS containing 1.25 µg/ml 4’,6-diamidino-2-phenylindole dihydrochloride (DAPI; ThermoFisher Scientific) and stored on ice until required. Immediately prior to sorting, cells were passed through a 35 µm filter. Single cells were sorted on a FACSAria II (BD Biosciences, San Jose, CA USA) using a 100 µm nozzle and a sheath pressure of 35 psi into individual wells of a 96-well plate each containing 200 µl Leibovitz’s L-15 medium with supplements as above including 20% (v/v) FBS. Cells were gated to select DAPI^low^ (live) single cell clones (Fig 1B). Fast-growing clonal cell lines confluent after three weeks of growth were expanded and confirmed to be mycoplasma-negative using the MycoAlert PLUS kit (Lonza, Basel, Switzerland) prior to freezing for long-term storage in liquid nitrogen. Clone Aag2-AF5 has been made available *via* ECACC (phe-culturecollections.org.uk; Public Health England, London, UK).

### Microscopy

Images were captured using an EVOS XL Core (Fig 1) or EVOS FL (Fig 5) Cell Imaging System (ThermoFisher Scientific). To measure transfection efficiency, images were captured 48 h after transient transfection with 300 ng pIEx-EGFP per well of a 12-well plate, each containing 1 × 10^6^ cells, at the time of cell seeding using TransIT-insect transfection reagent (Cambridge Biosciences, Cambridge, UK) as per manufacturer’s instructions. Transfection efficiency was calculated manually using Fiji (ImageJ) software (National Institutes of Health, Bethesda, MA USA) [69]. At least 900 individual cells were counted across three separate fields of view at 40X magnification for each experiment.

### PCR and RT-PCR

To analyse the genomic integration of PCLV, total DNA or total RNA was extracted from 1−3 × 10^6^ cells using the Quick-DNA or Quick-RNA Miniprep Kits (Zymo Research, Irvine, CA USA) respectively, as per manufacturer’s instructions. RNA samples were spiked with NDV prior to isolation. Nucleic acids were treated with DNase for 40 min at 37°C using the DNA-free DNA Removal Kit (ThermoFisher Scientific) or with RNase A (Sigma-Aldrich) for 1 h at 37°C as per manufacturers’ instructions. Nucleases were removed by re-purifying the nucleic acids as described above. cDNA was generated from RNA using the iScript cDNA Synthesis Kit (Bio-Rad, Hercules, CA USA) with random hexamers as per manufacturer’s instructions. PCR amplification was performed using the HOT FIREPol EvaGreen qPCR Supermix Plus (no ROX) (Solis BioDyne, Tartu, Estonia) at 95°C for 10 min followed by 26 cycles of 95°C for 15 s and 60°C for 30 s, as per manufacturer’s instructions. Primers (Sigma-Aldrich) were as follows; Rps7, prKM27F CCACGATCCCGCACTCTGA, prKM27R TACGCTTGCCGACGACTTCA; NDV, forward GACAATGCTTGATGGTGAAC, reverse CAATGCTGAGAGACAATAGGTC; PCLV L, prKM110F CACTGCTACACCGCCTAGAG, prKM110R TGACCTGTTGGCCTGTTGTT, prKM111F GCACCTTTAACAGGAGATGCAA, prKM111R ACTACGCCACAATGCGATGA, prKM112F GACTCCCCGATTGAGTAAAGAAC, prKM112R TCCAAGGAATCACTTTCTGATGC, prKM113F GTCGATTTCGAAGAAGTAGGTGC, prKM113R TCTATCGGTGATGTGCGTTCC, prKM231F AGGAGGCACAAATCAAGGTAGT, prKM231R GCGAGCTCACTTTGATGAATGG, prKM232F AGCCAGAGAAAGCAAACCAGA, prKM232R TCCATGTCATCAGTGTTGGTGT; PCLV M, prKM233F AGGCATGAAGACCTGGACTC, prKM233R GCATGCATCTGCTCTATGGG, prKM234F TTGCAGAGGAAGATCTCTGAGG, prKM234R TTCGCTTATCAGCCTGCAGTT, prKM235F GCCTGTCCCATCTGCGAAT, prKM235R AACCTGTGACTCGTGTGCAA, prKM236F AGCTGTTCTGGTAATGTTGTGGA, prKM236R TCTTCCAAGCAGGTTGGTTTG; PCLV S, prKM237F AGCAATAGATACGACTGCTAGTGA, prKM237R GCATTCATCTCCATACGCACA, prKM238F GCGTCATTCGTTTCGAGCAT, prKM238R TCAGCAGACGGAAATCGTTGT.

To test Aag2-derived clonal cell lines for the presence of insect-specific viruses, RNA was extracted from 1−3 × 10^6^ cells using the Quick-RNA Miniprep Kit (Zymo Research) as per manufacturer’s instructions. cDNA was synthesised using the Maxima H Minus First Strand cDNA Synthesis Kit (ThermoFisher Scientific) with random hexamers or gene-specific primers. PCR amplification was performed using the AccuPrime *Taq* High Fidelity DNA Polymerase (ThermoFisher Scientific) at 94°C for 2 min, followed by 35 cycles of 94°C for 30 s, 58°C for 30 s and 68°C for 1 min, followed by a final extension at 68°C for 5 min, as per manufacturer’s instructions. Primers were as follows; Rps7, prKM259F TGCTTTCGAGGGACAAATCGG, prKM259R AATTCGAACGTAACGTCACGTCC; CFAV, prKM258F TCATCTTATGTTGCACATGGACGC, prKM258R CACCCTCCGGAAATCCGATTG; PCLV L, prKM254F CATCAARRGATGAAGCCAGAGAAAG, prKM254R GTCTTTATGTTTTCTGTACAGCCATAAT; PCLV M, prKM256F AATGCAAACTGTTCTTGCAGATTCTG, prKM256R GTAGCTTAAAATCTGCGTCGTTAGT; PCLV S, prKM257F AATATAAATATTCAAACACCCCAGTTATAAG, prKM257R TCTGATCATTTAACATTCTCAGAGCTA.

### RT-qPCR

To measure PCLV levels, RNA was extracted from 1 × 10^6^ cells using 1 ml TRIzol reagent (ThermoFisher Scientific) and treated with DNase for 40 min at 37°C using the DNA-free DNA Removal Kit, as per manufacturers’ instructions. To measure immune gene induction, cells were seeded at 1 × 10^5^ cells/well in 96-well plates and stimulated one day later by replacing the culture medium with culture medium containing 1,000 colony-forming units (CFU)/cell heat-inactivated bacteria. RNA was isolated 24 h later using the Quick-RNA Miniprep Kit (Zymo Research) as per manufacturer’s instructions. RNA was reverse transcribed using the iScript cDNA Synthesis Kit with random hexamers as per manufacturer’s instructions. PCR amplification was performed using the HOT FIREPol EvaGreen qPCR Supermix Plus (no ROX) at 95°C for 10 min followed by 40 cycles of 95°C for 15 s and 60°C for 30 s, as per manufacturer’s instructions. Primers were as follows; PCLV L, prKM110F/R (see above); Rps7 (Genbank accession number XM_001660119), prKM27F CCACGATCCCGCACTCTGA, prKM27R TACGCTTGCCGACGACTTCA; Defensin D (XM_001657239), prKM14F TGCACCGGGGCCATTAC, prKM14R CAGGTGGCCCGTTTCAGG; Cecropin B (XM_001648640), prKM17F GAAGCTGGTCGGCTGAAGAA, prKM17R CAACGGGTAGTCCCTTCTGG; Cecropin D (XM_001649131), prKM16F AGCTGTTCGCAATTGTGCTGT, prKM16R TACAACAACCGGGAGAGCCTT.

### RNAi Assay

Cells were seeded at 1 × 10^5^ cells/well in 96-well plates and concurrently transiently transfected with 10 ng/well pKM50 (*Renilla*), 50 ng/well pKM19 (firefly luciferase) and 1 nM dsRNA directed against EGFP or firefly luciferase using TransIT-insect transfection reagent as per manufacturer’s instructions. Cells were harvested and luciferase activity measured two days post-transfection. Transfections for measuring transfection efficiency by firefly luciferase expression were set up in the same way, without the addition of dsRNA.

dsRNAs were generated by PCR amplification from plasmid templates pIEx-EGFP (EGFP dsRNA; primers prKM57F TAATACGACTCACTATAGGGCGTAAACGGCCACAAGTTCA, prKM57R TAATACGACTCACTATAGGGGGCGGACTTGAAGAAGTCGT) or pKM19 (firefly luciferase dsRNA; primers prKM168F TAATACGACTCACTATAGGGCAATCCGGAAGCGACCAACG, prKM168R TAATACGACTCACTATAGGGTTCCGCCCTTCTTGGCCTTT). Primers contain a 5’ T7 polymerase promoter used for *in vitro* transcription from gel purified amplicons using the MEGAshortscript *in vitro* transcription kit (ThermoFisher Scientific). dsRNA was gel purified prior to use.

### Bioinformatic Analysis of PCLV Insertions in Aag2 Genome

Our previously published full-length PCLV genome sequences from Aag2 cells (Genbank accession numbers KU936055, KU936056 and KU936057) [57] were searched against the Aag2 cell reference genome [70] using the BLAST function at vectorbase.org [71].

### Statistics

Heteroscedastic Student’s *t* test (assuming unequal variance) was performed in Microsoft Excel (Microsoft Corporation, Redmund, WA USA). For Fig 6, Student’s *t* test was calculated manually.

### Images

Graphs were plotted in Microsoft Excel. FACS images were generated in FlowJo (FlowJo LLC, Ashland, OR USA). Figures were prepared in Adobe Illustrator (Adobe Systems, San Jose, CA USA). Images were cropped, annotated and modified to optimise brightness and contrast only.

## RESULTS

### Establishment of Clonal Cell Lines Derived from Aag2 Cells

The *Ae. aegypti* Aag2 cell line forms a discontinuous monolayer of cells interspersed with three-dimensional cell clusters attached to the substrate and large rounded cells floating in isolation through the culture medium (Fig 1A). It is in principle possible that these morphological differences reflect underlying functional differences [53,54]. We therefore derived clonal cell lines from Aag2 cells to provide a more homogeneous and better-defined experimental background, for example for the generation of our previously reported CRISPR-edited cell lines [59,60]. Parental Aag2 cells were individually sorted into three 96-well plates using flow cytometry, with a stringent double gating scheme for single cells (Fig 1B). Of the 288 single cells plated, 90 clones expanded into multi-cell cultures in 24-well plates within three weeks. Fifteen of these clones were selected for further study and assigned reference numbers preceded by the prefix ‘AF’ (Aag2-AF1, Aag2-AF2 etc.). Clone Aag2-AF13 succumbed to fungal infection and is not discussed further. The set of clones was selected to be representative of the different morphologies observed across the population of clonal cell lines generated. Some clones resembled the parental Aag2 cell line (Fig 1Ci) while others did not form monolayers and instead either grew in large clusters attached to the substrate (Fig 1Cii) or grew floating individually and in aggregates in the culture medium (Fig 1Ciii). These clustered and floating morphologies are also observed in parental Aag2 cells (Fig 1A). Across all single-cell clones generated, these three morphologies (‘parental Aag2-like’, ‘clustered’, ‘rounded’) were represented at similar levels, with slightly more ‘clustered’ cell lines observed (Fig 1D). However, all of the clonal cell lines reverted back to the parental Aag2 morphology over time (Fig 1E), with this parental Aag2 morphology being stably maintained over many passages. This suggests that the different cell morphologies observed in the parental Aag2 cell line are not indicative of the presence of different cell types within the heterogeneous Aag2 cell culture.

We observed no gross differences in the growth kinetics of any of the Aag2-derived single-cell clones compared to the parental Aag2 cell line during routine culture. Before proceeding, we also confirmed that all clonal cell lines tested negative for mycoplasma. Note that we have used clone Aag2-AF5 as a well-defined single cell-derived starting point for the generation of Aag2 mutants using CRISPR [59,60].

### Parental Aag2 Cells Do Not Contain PCLV-Derived DNA Sequences Integrated into Their Genome

In mosquitoes and their derived cell lines, fragments of RNA virus genomes, including insect-specific virus genomes, can be reverse transcribed into DNA by endogenous reverse transcriptases [72,73], following which these fragments can become integrated into the cellular genome [74-77]. DNA sequences derived from members of the *Flaviviridae*, *Rhabdoviridae* and other viral families are known to be integrated within the Aag2 genome [54,75]. We were ultimately interested in studying the potential impact of persistent PCLV infection on acute superinfection with arboviruses in our Aag2-derived clonal cell lines, and therefore first investigated whether sequences derived from PCLV specifically are also integrated into the parental Aag2 cell genome. We performed a BLASTn search against the Aag2 reference genome [70] using our previously published full-length genome sequences for the PCLV known to infect the parental Aag2 cell line [57]. We did not identify any statistically significant (E-value <10^−5^) PCLV-derived sequences from any of the three viral genome segments (L, M, S) in the Aag2 reference genome sequence.

To rule out the possibility that fragments derived from persistent PCLV infection are integrated into the specific version of the parental Aag2 cell line growing in our lab, we designed primers to amplify short (50-150-nt) fragments covering each genome segment in 1,000-nt intervals. As a positive control we used RNA purified from parental Aag2 cells that had been subjected to a reverse transcription reaction (Fig 2A). To control for template that was only present in RNA form, we spiked the samples with the RNA virus Newcastle disease virus (NDV). To confirm that the RNA samples were not contaminated with residual DNA, we treated RNA with RNase or DNase prior to performing RT-PCR. Here, the mosquito genomic ribosomal subunit 7 (Rps7) served as a control, as this sequence is present in both RNA and DNA forms within the cell. For each PCLV-specific primer pair, and the NDV and Rps7 controls, RNase treatment eliminated the PCR signal, while DNase treatment did not (Fig 2A). This confirms the purity of the RNA samples. No PCLV signal was detected when PCR was performed on RNA that had not been subjected to reverse transcription, or on RNA isolated from the mammalian Madin-Darby canine kidney (MDCK) cell line, which should not contain genomic integrations of insect-specific virus sequences (Fig 2A). We then repeated the experiment using genomic DNA isolated from parental Aag2 cells, and detected no evidence of DNA sequences derived from PCLV (Fig 2A). Importantly, Rps7 was amplified when genomic DNA was treated with RNase, but not DNase, confirming the purity of the DNA samples (Fig 2A). Although we cannot exclude the possibility that the primers we designed missed smaller fragments of integrated PCLV sequence, our data suggest that PCLV fragments are not integrated into the genome of parental Aag2 cells, which is in agreement with data from other research groups [56].

**Fig 2.**
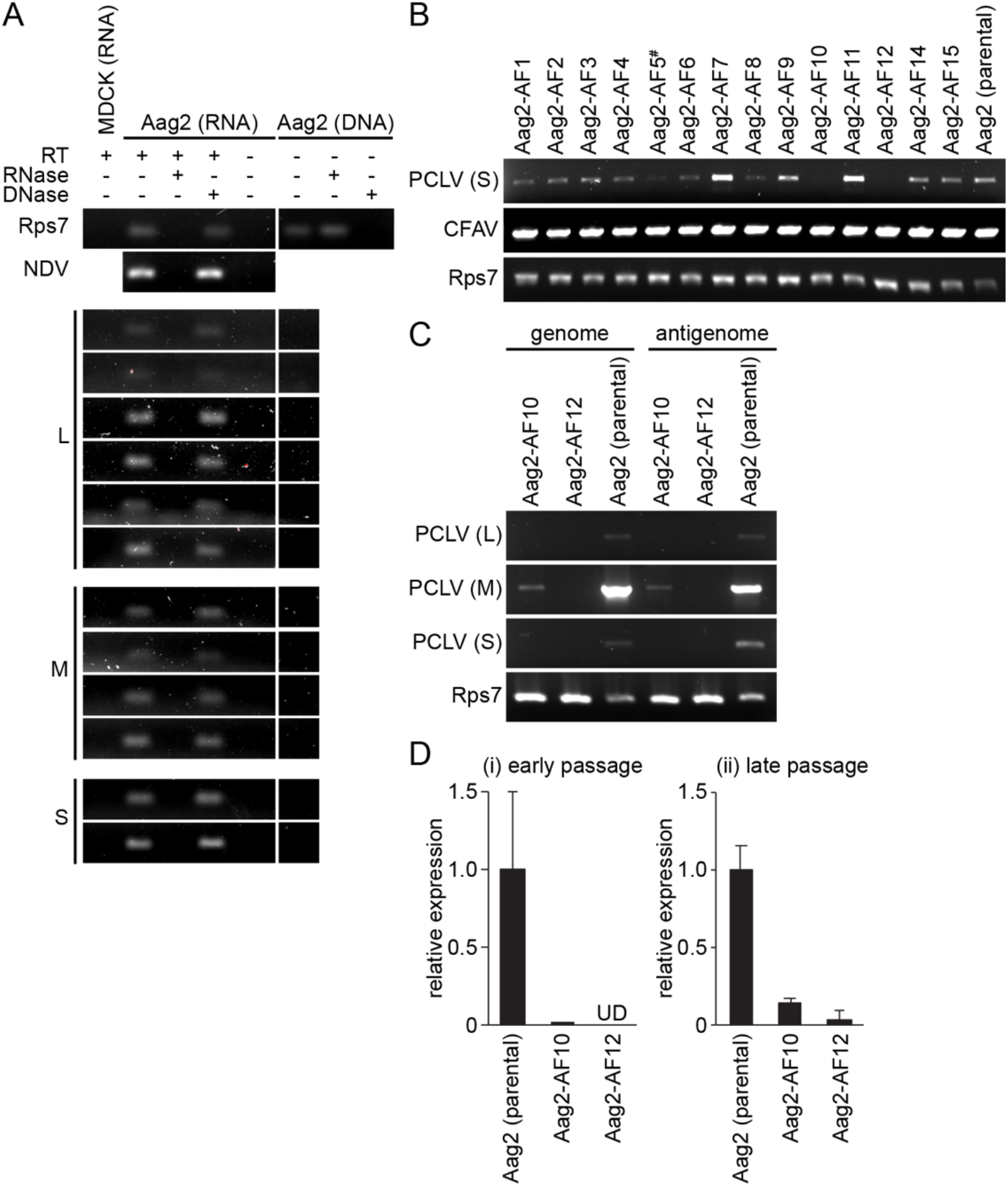
Aag2-AF10 and Aag2-AF12 cell lines harbour barely detectable levels of Phasi Charoen-like virus. (A) Short PCR amplicons spanning the three PCLV genome segments (L, M, S) amplified from RNA or genomic DNA isolated from parental Aag2 cells, either with or without a reverse transcription step (RT). Purified nucleic acids were treated with RNase or DNase prior to PCR. MDCK cell RNA, the cellular Rps7 gene/mRNA and the RNA virus NDV, which was spiked into cells immediately prior to RNA extraction, serve as controls. (B) Detection of the PCLV S segment and CFAV by RT-PCR in Aag2-derived clonal cell lines. Cellular Rps7 mRNA serves as a loading control. (C) Detection of the PCLV L, M and S genome (-ssRNA) and antigenome (+ssRNA) segments in select Aag2-derived clonal cell lines by sense-specific RT-PCR. Rps7 mRNA serves as a loading control. (D) PCLV L segment RT-qPCR ΔΔC_t_ (normalised to Rps7 mRNA) for select Aag2-derived clonal cell lines expressed relative to parental Aag2 cell line at (i) early passages (Aag2-AF10, passage 2; Aag2-AF12, passage 3) and (ii) later passages (Aag2-AF10, passage 8; Aag2-AF12, passage 12). Error bars represent standard deviation. UD, undetected. ^#^ Aag2-AF5 cell line used for CRISPR gene editing [59,60].

### Identification of Aag2-Derived Clonal Cell Lines Harbouring Consistently Low Levels of Persistent Insect-Specific Bunyavirus Infection

We next tested whether all of our Aag2-derived clonal cell lines still contained both of the insect-specific viruses known to persistently infect parental Aag2 cells. We detected CFAV RNA by RT-PCR in parental Aag2 cells and in all of our clonal cell lines, with cellular Rps7 RNA serving as a template control (Fig 2B). In contrast, clones Aag2-AF10 and Aag2-AF12 did not contain detectable levels of the PCLV S segment in this assay, while PCLV RNA was clearly detectable to varying degrees in parental Aag2 cells and in the other clones (Fig 2B). To verify this result, we performed strand-specific RT-PCR to amplify genome and antigenome sequences from each of the three PCLV genome segments, with Rps7 serving as a template control. In this experiment we did detect PCLV M segment RNA in the Aag2-AF10 clone at lower levels than the parental Aag2 cell line (Fig 2C). Although any amplification of the L and S segments were below the limit of detection, the presence of both genome and antigenome sequences for the M segment indicates that the virus must be replicating its RNA and therefore the L (RdRp) and S (nucleocapsid) segments, which are both required for genome replication, must also be present. No PCLV RNA was detected in the Aag2-AF12 clone in this assay.

Finally, we measured PCLV L segment RNA by RT-qPCR and detected low levels of PCLV RNA in the Aag2-AF10 clone, with PCLV RNA consistently maintained at lower levels relative to the parental Aag2 cell line over multiple cell passages (Fig 2D). We did not detect PCLV RNA at early passages in the Aag2-AF12 clone, though very low levels of PCLV were detected at later passages (Fig 2D). Overall, our data indicate that clones Aag2-AF10 and Aag2-AF12 harbour markedly reduced levels of PCLV infection that are maintained at consistently low levels over multiple cell passages.

### All Aag2-Derived Clonal Cell Lines Have a Functional RNAi Pathway

Derived cell lines can have markedly different characteristics compared to their parental cell lines, and several mosquito cell lines in particular are known to be immunodeficient compared to the cell lines they were derived from [43,53,54]. Before proceeding, we therefore tested our Aag2-derived clonal cell lines for RNAi functionality, since this immune pathway is defective in several mosquito cell lines, such as C6/36 and C7/10 [43,53,54]. In parental Aag2 cells, transient co-transfection of a constitutively active firefly luciferase reporter plasmid with a dsRNA directed against firefly luciferase significantly reduced luciferase expression compared to a non-specific dsRNA directed against GFP, confirming that RNAi is active in the parental Aag2 cell line (Fig 3). In contrast, the luciferase dsRNA did not significantly reduce reporter activity in RNAi-defective C6/36 cells (Fig 3). The luciferase-specific dsRNA significantly reduced firefly luciferase activity in all of the Aag2-derived clonal cell lines, confirming that all of the clones have a functional RNAi pathway (Fig 3). Furthermore, the clonal cell line Aag2-AF5 was previously shown to have a functional RNAi pathway as measured by the production of 21-nt siRNAs during viral infection [59,60].

**Fig 3.**
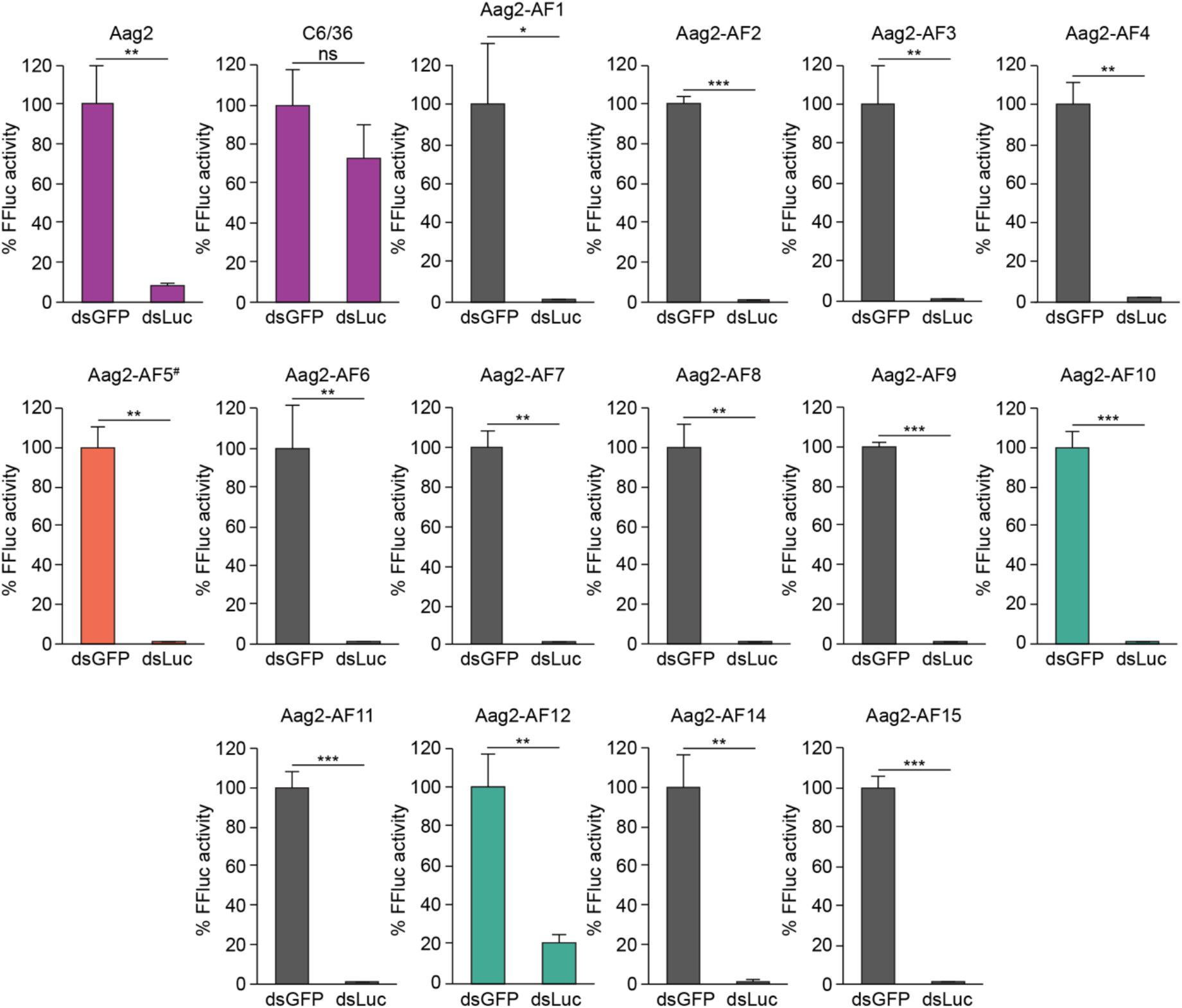
All Aag2-derived clonal cell lines have a functional RNAi pathway. C6/36 cells, the parental Aag2 cell line and its derived clonal cell lines were transiently transfected with plasmids constitutively expressing firefly luciferase and *Renilla* luciferase (transfection control) in the presence of dsRNA directed against GFP (dsGFP) or firefly luciferase (dsLuc). Mean *Renilla*-normalised firefly luciferase (FFluc) expression is expressed relative to the dsGFP negative control. * *P* < 0.05; ** *P* < 0.01; *** *P* < 0.001; ns, not significant (one-tailed Student’s *t* test). Error bars represent standard deviation. ^#^ Aag2-AF5 cell line used for CRISPR gene editing [59,60] is highlighted in orange. PCLV-low clones Aag2-AF10 and Aag2-AF12 are highlighted in green; parental Aag2 cells and C6/36 cells (negative control) are shown in purple.

### Pre-Existing Persistent Infection With PCLV Does Not Modulate Acute Superinfection With Flaviviruses

The isolation of the clonal cell lines Aag2-AF10 and Aag2-AF12 provided an opportunity to test whether a pre-existing persistent infection with an insect-specific virus (in this case PCLV) modulates the replication of arboviruses in cell culture by comparing these PCLV-low clones to clones harbouring higher levels of PCLV. We started by testing the flavivirus DENV serotype 2 (DENV-2), which replicated with a peak in titres six days post-infection in a one-step growth curve at high multiplicity of infection (MOI 2) in the parental Aag2 cell line (Fig 4A). DENV-2 replicated with similar kinetics (1, 2, 3 days post-infection) and to a similar level three days post-infection in clones Aag2-AF10 and Aag2-AF12 relative to the parental Aag2 cell line at MOI 2 (Fig 4B). This indicates that the markedly suppressed PCLV infection in clones Aag2-AF10 and Aag2-AF12 had very little impact on DENV-2 replication. Although DENV-2 exhibited somewhat different growth kinetics across the Aag2-derived clonal cell lines, titres three days post-infection did not deviate from the parental Aag2 cell line by more than one log, and were close to within half a log of the parental Aag2 cell line for all clones (Fig 4B).

**Fig 4.**
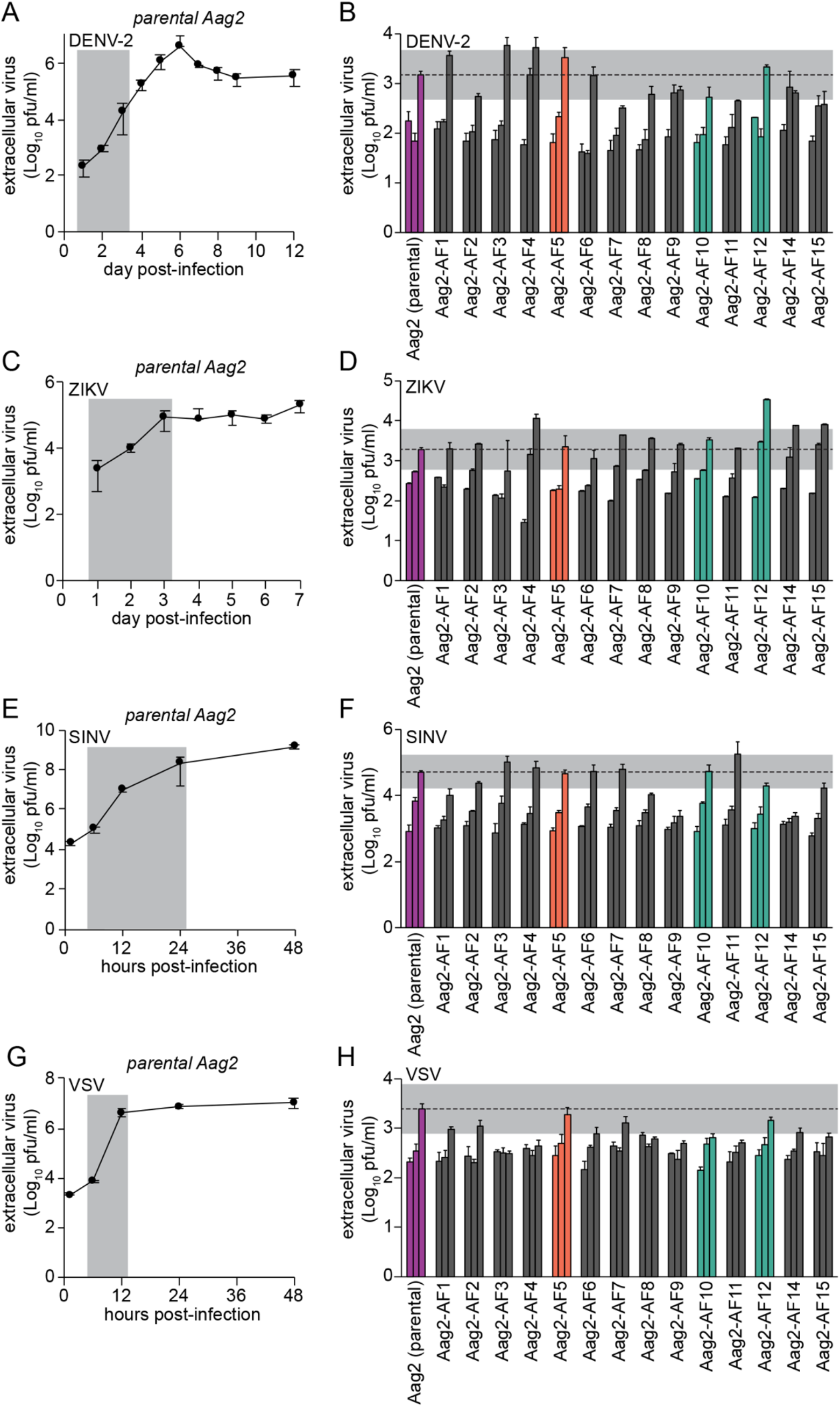
Susceptibility of Aag2-derived clonal cell lines to infection with arboviruses. (A, C, E, G) Single-step growth kinetics of DENV-2 (A), ZIKV (C), SINV (E) and VSV (G) in the parental Aag2 cell line (MOI 2). Grey shading highlights time points tested in Aag2-derived clonal cell lines. (B, D, F, H) Replication of DENV-2 (B) and ZIKV (D) at 1, 2 and 3 days post-infection, replication of SINV (F) at 6, 12 and 24 hours post-infection (hpi), and replication of VSV (H) at 6, 9 and 12 hpi in the parental Aag2 cell line and its derived clonal cell lines (all MOI 2). Grey shading indicates 0.5 Log_10_ above and 0.5 Log_10_ below peak extracellular titres detected in parental Aag2 cell line. Error bars represent standard deviation. Parental Aag2 cell line highlighted in purple; Aag2-AF5 cell line used for CRISPR gene editing [59,60] highlighted in orange; Aag2-AF10 and Aag2-AF12 cell lines with low PCLV levels highlighted in green.

We next tested the flavivirus ZIKV, which replicated with a peak in titres three days post-infection in the parental Aag2 cell line at MOI 2 (Fig 4C). While ZIKV replicated with faster kinetics (1, 2, 3 days post-infection) and to a more than one-log higher titre at its peak in clone Aag2-AF12, the growth kinetics and peak titres were similar to the parental Aag2 cell line in clone Aag2-AF10 (Fig 4D). Therefore, although clone Aag2-AF12 appears to be more permissive to ZIKV replication, this is not linked to PCLV levels, which are also reduced in clone Aag2-AF10. Again, ZIKV replication kinetics varied somewhat across the other clones, but fell close to within half a log from the parental Aag2 cell line at their peak.

Replication kinetics of both DENV-2 and ZIKV in clone Aag2-AF5 were comparable to the parental Aag2 cell line (Fig 4B and 4D).

### Pre-Existing Persistent Infection With PCLV Does Not Modulate Acute Superinfection With the Alphavirus Sindbis Virus

Alphaviruses, like flaviviruses, are +ssRNA viruses, and we next tested replication of the model alphavirus SINV in our Aag2-derived clonal cell lines. SINV replication peaked 48 hpi in parental Aag2 cells infected at MOI 2 (Fig 4E). SINV replication kinetics (6, 12, 24 hpi) and peak viral titres were similar in clones Aag2-AF10 and Aag2-AF12 compared to parental Aag2 cells (Fig 4F), indicating that persistent PCLV infection does not markedly alter SINV replication. Again, variability in the replication kinetics and peak titres of SINV were observed across the other Aag2-derived clonal cell lines. SINV replication kinetics and peak titres in clone Aag2-AF5 were comparable to the parental Aag2 cell line.

### Pre-Existing Persistent Infection With PCLV Does Not Modulate Acute Superinfection With the Rhabdovirus Vesicular Stomatitis Virus

As a contrast to the +ssRNA arboviruses tested, we next tested the −ssRNA rhabdovirus VSV. In a one-step growth curve (MOI 2), VSV replication peaked 12 hpi in parental Aag2 cells (Fig 4G). As for the other viruses tested, peak titres of VSV in clones Aag2-AF10 and Aag2-AF12 were within one log compared to the parental Aag2 cell line at 12 hpi, with some variability in replication kinetics (6, 9, 12 hpi) observed across all Aag2-derived single-cell clones (Fig 4H). VSV replicated similarly in clone Aag2-AF5 and the parental Aag2 cell line.

Overall, we therefore conclude that pre-existing persistent infection with PCLV does not notably alter the replication of a diverse range of +ssRNA and −ssRNA arboviruses, since replication kinetics and peak titres of DENV-2, ZIKV, SINV and VSV were not markedly different from the parental Aag2 cell line in clones Aag2-AF10 and Aag2-AF12, which harbour drastically reduced levels of persistent PCLV infection.

### Transfection Efficiency of Clone Aag2-AF5

Next, we further characterised clone Aag2-AF5 with the goal of providing a better-defined Aag2-derived cell line for the research community. Clone Aag2-AF5 was selected because, of all the isolated clones, arboviral infectivity in Aag2-AF5 cells was most similar to the parental Aag2 cells (Fig 4). Furthermore, this clone formed a more uniform monolayer and was more resilient and easier to handle in culture than parental Aag2 cells.

First, we tested the transfection efficiency of this clone by transient transfection with a constitutive GFP expression plasmid (Fig 5A). A similar proportion of Aag2-AF5 cells (56%) were detectably GFP-positive compared to the parental Aag2 cell line (47%) (Fig 5B); differences were not statistically significant. However, the GFP signal was brighter in Aag2-AF5 cells (Fig 5A). This higher level of transgene expression in Aag2-AF5 cells was confirmed by transient transfection with a constitutively active firefly luciferase reporter plasmid (Fig 5C). Therefore, while the overall proportion of cells transiently expressing a transgene is comparable for clone Aag2-AF5 and the parental Aag2 cell line, individual Aag2-AF5 cells express transgenes to higher levels, making this clone well-suited for molecular experiments including gene editing using CRISPR [59,60].

**Fig 5.**
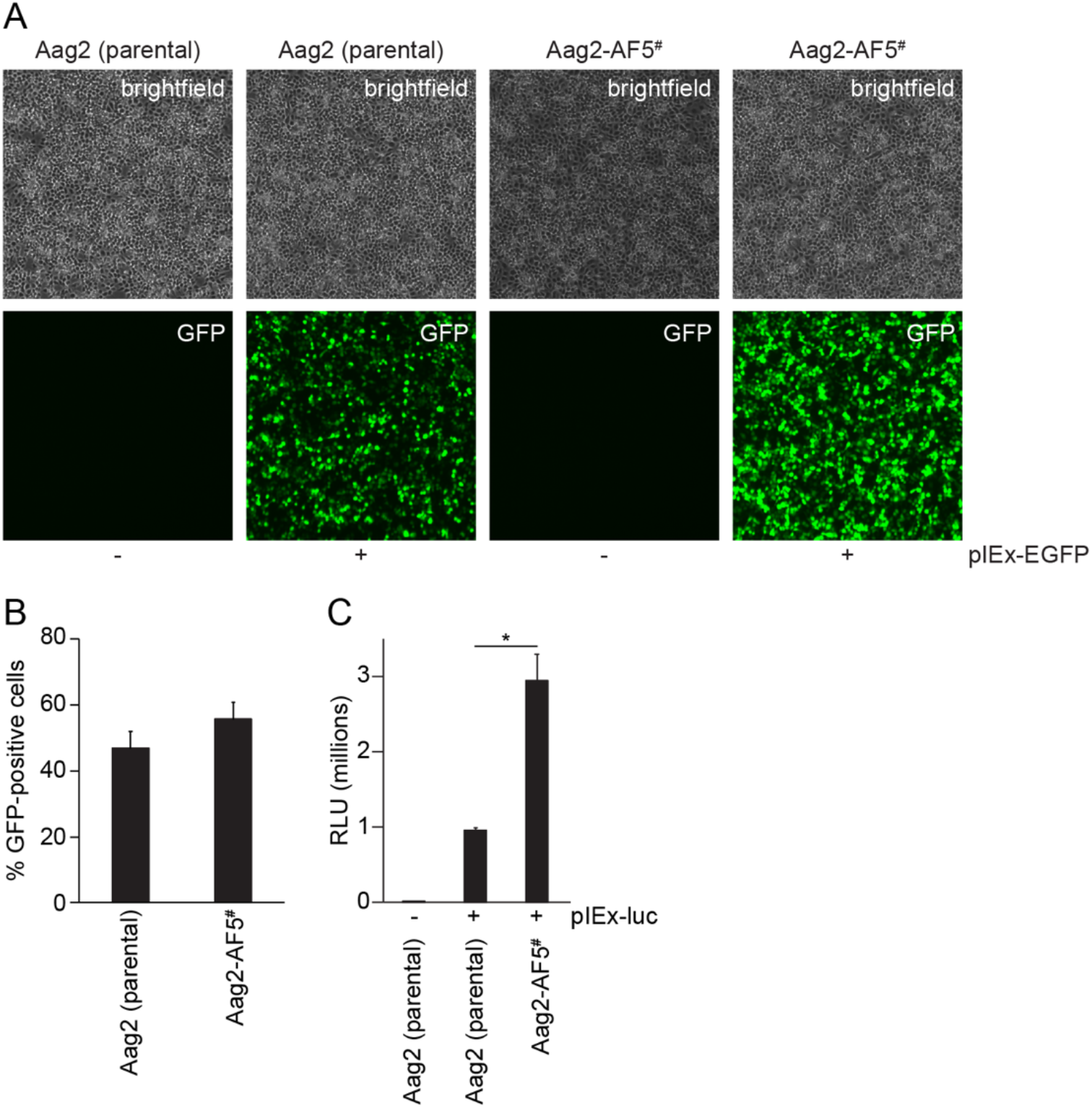
Transfection efficiency of clone Aag2-AF5 relative to parental Aag2 cells. (A) Cells were imaged at 10X magnification 48 h after transient transfection with a constitutively active GFP expression vector (pIEx-EGFP). (B) Quantification of transfection efficiency; differences are non-significant. (C) Cells were transiently transfected with a constitutively active firefly luciferase reporter plasmid (pIEx-luc) and luciferase activity was measured four days later. * *P* < 0.05 (two-tailed Student’s *t* test). RLU, relative light units. All error bars represent standard deviation. ^#^ Aag2-AF5 cell line used for CRISPR gene editing [59,60].

### Antimicrobial Peptide Induction in Clone Aag2-AF5

Clone Aag2-AF5 was already confirmed to have an active antiviral RNAi pathway (Fig 3) [59,60], and we next tested whether this clone was also competent for antimicrobial peptide induction *via* inducible innate immune signalling pathways. When stimulated with heat-inactivated Gram-negative (*Escherichia coli*) or Gram-positive (*Listeria monocytogenes* or *Staphylococcus aureus*) bacteria, which are well-defined stimuli of inducible innate immune signalling pathways [51], upregulation of the antimicrobial peptides defensin D (DefD), cecropin B (CecB) and cecropin D (CecD) was detected in both parental Aag2 cells and clone Aag2-AF5 with all stimuli (Fig 6). There was however some variability in the relative levels of antimicrobial peptide induction in clone Aag2-AF5 compared to the parental cell line for different gene/stimulus combinations. Thus, all tested antimicrobial peptide genes were less inducible in clone Aag2-AF5 with *E. coli* stimulation (Fig 6A), and CecD was also less inducible in clone Aag2-AF5 for all stimuli tested (Fig 6Aiii, 6Biii and 6Ciii), though some of these differences were not significant. In contrast, DefD and CecB were more inducible in clone Aag2-AF5 during stimulation with Gram-positive bacteria compared to the parental Aag2 cell line (Fig 6Bi, 6Bii, 6Ci and 6Cii), with some of these differences again being non-significant. Therefore, while there may be subtle differences in the immune sensitivity of clone Aag2-AF5 in terms of antimicrobial peptide production compared to parental Aag2 cells, there is no gross defect in inducible innate immune pathways in response to bacterial stimulation.

**Fig 6.**
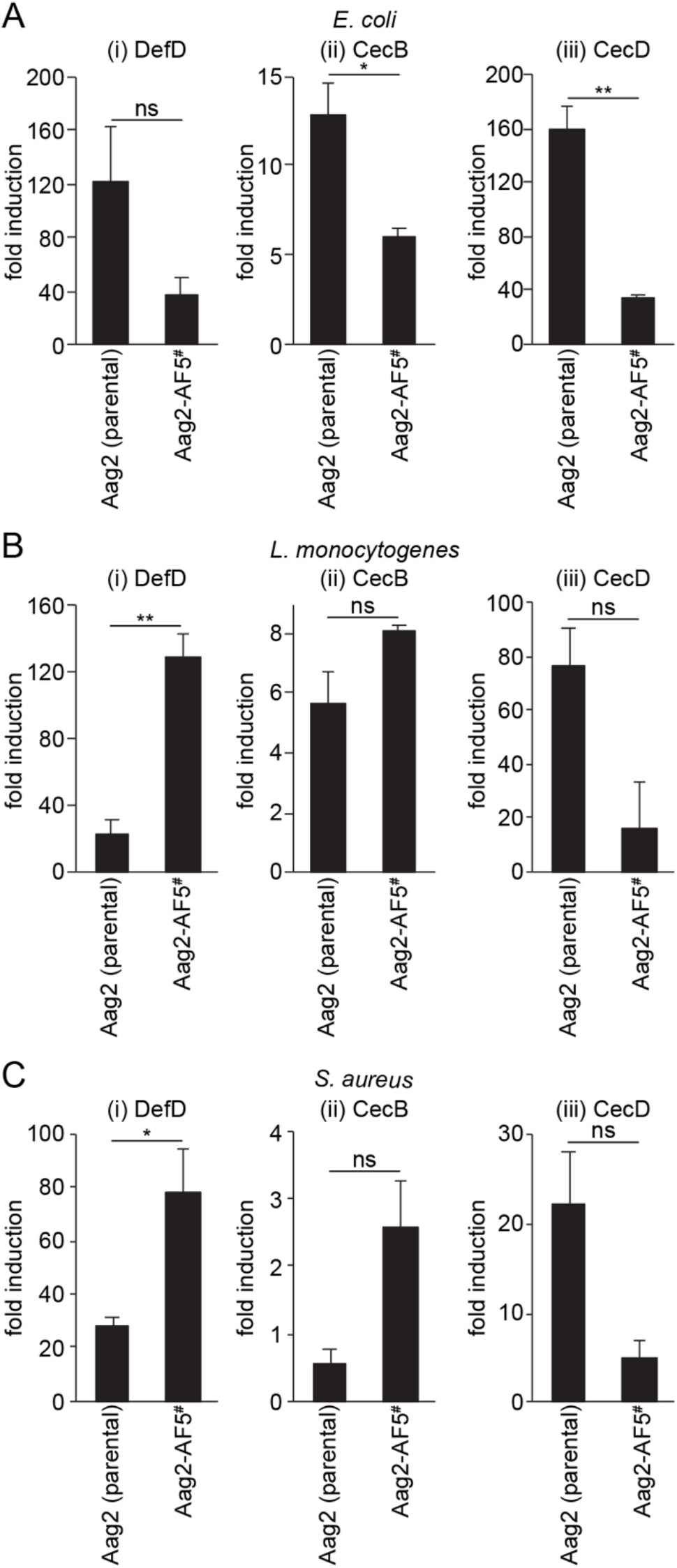
Antimicrobial peptide induction in clone Aag2-AF5 compared to parental Aag2 cells. Cells were stimulated with heat-inactivated *E. coli* (A), *L. monocytogenes* (B) or *S. aureus* (C) for 24 h and induction of DefD (i), CecB (ii) or CecD (iii) was measured by RT-qPCR. Gene induction is relative to the respective unstimulated cell line. * *P* < 0.05; ** *P* < 0.01; ns, not significant (two-tailed Student’s *t* test). Error bars represent standard error of the mean. ^#^ Aag2-AF5 cell line used for CRISPR gene editing [59,60].

## DISCUSSION

In this study, we generated clonal cell lines from the widely used Aag2 cell line. While the different morphologies within the parental Aag2 cell line could have been indicative of the presence of different embryo-derived cell types, this appears not to be the case as all of the clonal cell lines reverted back to the parental Aag2 cell morphology, with cell clusters and floating cells above the monolayer. In our hands, the Aag2 morphology is highly susceptible to culture conditions, and therefore the different cell morphologies may instead reflect cellular responses to growth phase, cell density or nutrient status.

Across all of the clonal Aag2-derived cell lines, there was minor variability in the replication kinetics and peak titres for all of the viruses tested. These effects were not consistent across viruses or cell lines and did not overall correlate consistently with the presence or level of insect-specific viruses in the culture. Neither are these effects likely to be linked to variability in immune responses because all clonal cell lines had a functional RNAi pathway, considered to be the major antiviral immune pathway in insects [49,50]. However, we did not extensively test inducible immune pathways such as Toll, IMD or Jak-STAT signalling in all clones. The clonal cell lines may exhibit variability in the expression of pro- or anti-viral factors, and could therefore be useful for identifying viral restriction factors or host proteins required for viral replication in mosquito cells.

### Clone Aag2-AF5 as a Defined Cell Line for the Arbovirus Research Community

There has been a drive to improve data reproducibility, with the standardisation of experimental methods and tools representing one important means of achieving this end [78]. We believe that our Aag2-derived clone Aag2-AF5 represents a useful standardised *Ae. aegypti* cell line for the arbovirus research community, and have made the cell line available via ECACC (phe-culturecollections.org.uk). The parental Aag2 cell line has previously been shown to be a valuable tool for studying mosquito immune responses to arbovirus infection [49,50], and we confirmed that Aag2-AF5 cells are also competent for RNAi and antimicrobial peptide induction. Furthermore, the viruses tested all replicated with similar kinetics and to similar peak titres in Aag2-AF5 cells compared to the parental Aag2 cell line, and this clone was therefore chosen to allow comparison to experiments performed in parental Aag2 cells. Morphologically, Aag2-AF5 cells are similar to the parental Aag2 cell line, and easier to work with in culture. Both CFAV and PCLV remain present in Aag2-AF5 cells, though PCLV is present at slightly reduced levels compared to the parental Aag2 cell line (Fig 2B). Aag2-AF5 cells are readily transfected and express exogenous proteins to high levels. We have also previously shown that Aag2-AF5 cells are easily gene edited using CRISPR [59], and Aag2-AF5 cells provide a more homogeneous background for gene editing experiments compared to the parental Aag2 cell line.

We have shared Aag2-AF5 cells widely within the research community, and their availability via ECACC should further increase their utility in standardising cell culture experiments to provide more reproducible data on arbovirus-vector interactions in *Ae. aegypti* cells.

### Impact of Persistent Insect-Specific Virus Infection on Arbovirus Replication

Two of the clonal Aag2-derived cell lines (Aag2-AF10 and Aag2-AF12) exhibited markedly reduced levels of persistent PCLV infection. PCLV levels remained consistently low over multiple passages, and in the case of Aag2-AF12 cells were so low as to be only intermittently detectable. In the wild, insect-specific viruses primarily cause persistent infection of mosquitoes, and comparison of the PCLV-low clones Aag2-AF10 and Aag2-AF12 to parental Aag2 cells allowed us to more accurately model these natural conditions than previous studies that tested the impact of acute insect-specific virus infection on arbovirus replication [31,34-37,41,42].

We observed no consistent impact of PCLV on the replication of representative flaviviruses (DENV-2, ZIKV), alphaviruses (SINV) or rhabdoviruses (VSV), representing both +ssRNA and −ssRNA arboviruses. To our knowledge, only one previous study tested the impact of persistent insect-specific virus (AeAV) infection on arbovirus (DENV) replication in immunocompetent *Aedes spp.* cell cultures [19], with at most a minor reduction in DENV replication in the presence of AeAV. Our data agree with this study in that no major impact of persistent insect-specific virus infection was observed on arbovirus replication, with our study extending this observation to a broader set of arboviral families. A study by Kuwata *et al.* also found that persistent insect-specific virus (CxFV) infection did not reduce, and in fact increased, arbovirus (DENV, JEV) replication in a *Culex tritaeniorhynchus* cell line [29]. To our knowledge there are no studies testing the impact of persistent insect-specific virus infection on arbovirus replication *in vivo* in *Aedes spp.*, however our cell line data are also in agreement with an *in vivo* study in *Culex spp.* that found no impact of persistent insect-specific virus (CxFV) infection on arbovirus (West Nile virus, WNV) transmission [32].

The lack of impact of persistent insect-specific virus infection on arbovirus replication in *Aedes spp.* cell culture is noteworthy because a number of studies have reported reduced replication of arboviruses in the presence of acute insect-specific virus infection in cell lines [31,34-37] and in the presence of acute insect-specific virus infection *in vivo* [31,35-37] in both *Aedes spp.* and *Culex spp*. Arboviruses have been shown to be affected differently when entering cells harbouring acute *versus* persistent insect-specific virus infection [40], which likely expose arboviruses to markedly different cellular environments. For instance, immune responses likely differ under acute *versus* persistent infection scenarios, and persistent infection with at least one insect-specific virus (Culex Y virus) has been shown to modulate RNAi responses [79]. Therefore, studies linking acute or persistent insect-specific virus infection to reduced arbovirus replication in the immunocompromised C6/36 cell line [32,33,39,40] may also not fully reflect the effects of persistent insect-specific virus infection in natural settings.

Our findings and those of others [19,32] suggest that the persistent insect-specific virus infections encountered by arboviruses in nature and in mosquitoes hypothetically infected for environmental release may not reduce, and may in fact enhance [29], arbovirus transmission. However, further studies are required to reconcile the contradictory observations made by different research groups, which may be influenced by insect-specific virus infection status (acute *versus* persistent), as well as the tripartite combination of arbovirus, insect-specific virus and mosquito species. Representative *in vivo* studies in particular are much needed, since cell culture experiments do not fully recapitulate all facets of the arbovirus infection process in mosquitoes.

We therefore provide new insights that may have important implications for the use of insect-specific viruses as biocontrol agents to reduce the transmission of arboviruses. Furthermore, clone Aag2-AF5 represents a valuable new clonal and better-defined cell line to provide a more standardised system for studying arbovirus-vector interactions in cell culture.

## ACKNOWLEDGEMENTS

None.

## AUTHORS’ CONTRIBUTIONS

KM and ADD conceived the project. KM, AF-S and ADD secured funding. ACF, LEW, TAR and KM performed experiments. All authors were involved in the analysis and interpretation of data. KM wrote the manuscript. All authors contributed to the preparation of the manuscript. All authors read and approved the final manuscript.

## REFERENCES

1. Weaver SC, Charlier C, Vasilakis N, Lecuit M. Zika, Chikungunya, and Other Emerging Vector-Borne Viral Diseases. Annu Rev Med. 2017;69: 395–408. doi:10.1146/annurev-med-050715-105122

2. Bolling BG, Weaver SC, Tesh RB, Vasilakis N. Insect-Specific Virus Discovery: Significance for the Arbovirus Community. Viruses. Multidisciplinary Digital Publishing Institute; 2015;7: 4911–4928. doi:10.3390/v7092851

3. Cook S, Moureau G, Kitchen A, Gould EA, de Lamballerie X, Holmes EC, et al. Molecular evolution of the insect-specific flaviviruses. J Gen Virol. 2012;93: 223–234. doi:10.1099/vir.0.036525-0

4. Kuno G, Chang G-JJ. Biological transmission of arboviruses: reexamination of and new insights into components, mechanisms, and unique traits as well as their evolutionary trends. Clinical Microbiology Reviews. American Society for Microbiology Journals; 2005;18: 608–637. doi:10.1128/CMR.18.4.608-637.2005

5. Weaver SC, Barrett ADT. Transmission cycles, host range, evolution and emergence of arboviral disease. Nat Rev Micro. 2004;2: 789–801. doi:10.1038/nrmicro1006

6. Kramer LD. Complexity of virus-vector interactions. Current Opinion in Virology. 2016;21: 81–86. doi:10.1016/j.coviro.2016.08.008

7. Souza-Neto JA, Powell JR, Bonizzoni M. Aedes aegypti vector competence studies: A review. Infect Genet Evol. 2018;67: 191–209. doi:10.1016/j.meegid.2018.11.009

8. Beerntsen BT, James AA, Christensen BM. Genetics of Mosquito Vector Competence. Microbiology and Molecular Biology Reviews. American Society for Microbiology; 2000;64: 115–137. doi:10.1128/MMBR.64.1.115-137.2000

9. Kramer MC, Liang D, Tatomer DC, Gold B, March ZM, Cherry S, et al. Combinatorial control of Drosophila circular RNA expression by intronic repeats, hnRNPs, and SR proteins. Genes Dev. Cold Spring Harbor Lab; 2015;29: 2168–2182. doi:10.1101/gad.270421.115

10. Walker T, Johnson PH, Moreira LA, Iturbe-Ormaetxe I, Frentiu FD, McMeniman CJ, et al. The wMel Wolbachia strain blocks dengue and invades caged Aedes aegypti populations. Nature. Nature Publishing Group; 2011;476: 450–453. doi:10.1038/nature10355

11. Moreira LA, Iturbe-Ormaetxe I, Jeffery JA, Lu G, Pyke AT, Hedges LM, et al. A Wolbachia Symbiont in Aedes aegypti Limits Infection with Dengue, Chikungunya, and Plasmodium. Cell. Elsevier Ltd; 2009;139: 1268–1278. doi:10.1016/j.cell.2009.11.042

12. Blagrove MSC, Arias-Goeta C, Failloux AB, Sinkins SP. Wolbachia strain wMel induces cytoplasmic incompatibility and blocks dengue transmission in Aedes albopictus. Proc Natl Acad Sci. 2012;109: 255–260. doi:10.1073/pnas.1112021108

13. Bian G, Xu Y, Lu P, Xie Y, Xi Z. The Endosymbiotic Bacterium Wolbachia Induces Resistance to Dengue Virus in Aedes aegypti. Schneider DS, editor. PLoS Pathog. 2010;6: e1000833. doi:10.1371/journal.ppat.1000833.s001

14. Ferguson NM. Challenges and opportunities in controlling mosquito-borne infections. Nature. Nature Publishing Group; 2018;559: 490–497. doi:10.1038/s41586-018-0318-5

15. Öhlund P, Lundén H, Blomström A-L. Insect-specific virus evolution and potential effects on vector competence. Virus Genes. Springer US; 2019;64: 1–11. doi:10.1007/s11262-018-01629-9

16. Zakrzewski M, Rasic G, Darbro J, Krause L, Poo YS, Filipovic I, et al. Mapping the virome in wild-caught Aedes aegypti from Cairns and Bangkok. Sci Rep. Nature Publishing Group; 2018;8: 4690. doi:10.1038/s41598-018-22945-y

17. Zhang X, Huang S, Jin T, Lin P, Huang Y, Wu C, et al. Discovery and high prevalence of Phasi Charoen-like virus in field-captured Aedes aegypti in South China. Virology. 2018;523: 35–40. doi:10.1016/j.virol.2018.07.021

18. Yamao T, Eshita Y, Kihara Y, Satho T, Kuroda M, Sekizuka T, et al. Novel virus discovery in field-collected mosquito larvae using an improved system for rapid determination of viral RNA sequences (RDV ver4.0). Archives of Virology. 2009;154: 153–158. doi:10.1007/s00705-008-0285-5

19. Parry R, Asgari S. Aedes anphevirus (AeAV): an insect-specific virus distributed worldwide in Aedes aegypti mosquitoes that has complex interplays with Wolbachia and dengue virus infection in cells. J Virol. American Society for Microbiology; 2018;: JVI.00224-18. doi:10.1128/JVI.00224-18

20. Vasilakis N, Guzman H, Firth C, Forrester NL, Widen SG, Wood TG, et al. Mesoniviruses are mosquito-specific viruses with extensive geographic distribution and host range. Virol J. 2014;11: 97. doi:10.1186/1743-422X-11-97

21. Huhtamo E, Putkuri N, Kurkela S, Manni T, Vaheri A, Vapalahti O, et al. Characterization of a novel flavivirus from mosquitoes in northern europe that is related to mosquito-borne flaviviruses of the tropics. J Virol. 2009;83: 9532–9540. doi:10.1128/JVI.00529-09

22. Sangdee K, Pattanakitsakul S-N. Comparison of Mosquito Densoviruses: Two Clades of Viruses Isolated from Indigenous Mosquitoes. Southeast Asian Journal of Tropical Medicine and Public Health. 2013;44:586–593.

23. Junglen S, Kopp A, Kurth A, Pauli G, Ellerbrok H, Leendertz FH. A new flavivirus and a new vector: characterization of a novel flavivirus isolated from uranotaenia mosquitoes from a tropical rain forest. J Virol. 2009;83: 4462–4468. doi:10.1128/JVI.00014-09

24. Nasar F, Palacios G, Gorchakov RV, Guzman H, Da Rosa APT, Savji N, et al. Eilat virus, a unique alphavirus with host range restricted to insects by RNA replication. Proc Natl Acad Sci. 2012;109: 14622–14627. doi:10.1073/pnas.1204787109

25. Cook S, Bennett SN, Holmes EC, De Chesse R, Moureau G, de Lamballerie X. Isolation of a new strain of the flavivirus cell fusing agent virus in a natural mosquito population from Puerto Rico. J Gen Virol. 2006;87: 735–748. doi:10.1099/vir.0.81475-0

26. Cook S, Moureau G, Harbach RE, Mukwaya L, Goodger K, Ssenfuka F, et al. Isolation of a novel species of flavivirus and a new strain of Culex flavivirus (Flaviviridae) from a natural mosquito population in Uganda. J Gen Virol. 2009;90: 2669–2678. doi:10.1099/vir.0.014183-0

27. Newman CM, Cerutti F, Anderson TK, Hamer GL, Walker ED, Kitron UD, et al. Culex Flavivirus and West Nile Virus Mosquito Coinfection and Positive Ecological Association in Chicago, United States. Vector-Borne Zoonotic Dis. 2011;11: 1099–1105. doi:10.1089/vbz.2010.0144

28. Calzolari M, Ze-Ze L, Ruzek D, Vazquez A, Jeffries C, Defilippo F, et al. Detection of mosquito-only flaviviruses in Europe. J Gen Virol. 2012;93: 1215–1225. doi:10.1099/vir.0.040485-0

29. Kuwata R, Isawa H, Hoshino K, Sasaki T, Kobayashi M, Maeda K, et al. Analysis of Mosquito-Borne Flavivirus Superinfection in Culex tritaeniorhynchus (Diptera: Culicidae) Cells Persistently Infected with Culex Flavivirus (Flaviviridae). Journal of Medical Entomology. 2015;52: 222–229. doi:10.1093/jme/tju059

30. Zhang G, Asad S, Khromykh AA, Asgari S. Cell fusing agent virus and dengue virus mutually interact in Aedes aegypti cell lines. Sci Rep. Nature Publishing Group; 2017;7: 6935. doi:10.1038/s41598-017-07279-5

31. Romo H, Kenney JL, Blitvich BJ, Brault AC. Restriction of Zika virus infection and transmission in Aedes aegypti mediated by an insect-specific flavivirus. Emerg Microbes Infect. Nature Publishing Group; 2018;7: 181. doi:10.1038/s41426-018-0180-4

32. Bolling BG, Olea-Popelka FJ, Eisen L, Moore CG, Blair CD. Transmission dynamics of an insect-specific flavivirus in a naturally infected Culex pipiens laboratory colony and effects of co-infection on vector competence for West Nile virus. Virology. Elsevier Inc; 2012;427: 90–97. doi:10.1016/j.virol.2012.02.016

33. Hobson-Peters J, Yam AWY, Lu JWF, Setoh YX, May FJ, Kurucz N, et al. A New Insect-Specific Flavivirus from Northern Australia Suppresses Replication of West Nile Virus and Murray Valley Encephalitis Virus in Co-infected Mosquito Cells. Wang T, editor. PLoS One. 2013;8: e56534. doi:10.1371/journal.pone.0056534.s001

34. Kenney JL, Solberg OD, Langevin SA, Brault AC. Characterization of a novel insect-specific flavivirus from Brazil: potential for inhibition of infection of arthropod cells with medically important flaviviruses. J Gen Virol. 2014;95: 2796–2808. doi:10.1099/vir.0.068031-0

35. Goenaga S, Kenney JL, Duggal NK, Delorey M, Ebel GD, Zhang B, et al. Potential for Co-Infection of a Mosquito-Specific Flavivirus, Nhumirim Virus, to Block West Nile Virus Transmission in Mosquitoes. Viruses. Multidisciplinary Digital Publishing Institute; 2015;7: 5801–5812. doi:10.3390/v7112911

36. Nasar F, Erasmus JH, Haddow AD, Tesh RB, Weaver SC. Eilat virus induces both homologous and heterologous interference. Virology. Academic Press; 2015;484: 51–58. doi:10.1016/j.virol.2015.05.009

37. Hall-Mendelin S, McLean BJ, Bielefeldt-Ohmann H, Hobson-Peters J, Hall RA, van den Hurk AF. The insect-specific Palm Creek virus modulates West Nile virus infection in and transmission by Australian mosquitoes. Parasit Vectors. BioMed Central; 2016;9: 414. doi:10.1186/s13071-016-1683-2

38. Schultz MJ, Frydman HM, Connor JH. Dual Insect specific virus infection limits Arbovirus replication in Aedes mosquito cells. Virology. 2018;518: 406–413. doi:10.1016/j.virol.2018.03.022

39. Fujita R, Kato F, Kobayashi D, Murota K, Takasaki T, Tajima S, et al. Persistent viruses in mosquito cultured cell line suppress multiplication of flaviviruses. Heliyon. 2018;4: e00736. doi:10.1016/j.heliyon.2018.e00736

40. Burivong P, Pattanakitsakul S-N, Thongrungkiat S, Malasit P, Flegel TW. Markedly reduced severity of Dengue virus infection in mosquito cell cultures persistently infected with Aedes albopictus densovirus (AalDNV). Virology. 2004;329: 261–269. doi:10.1016/j.virol.2004.08.032

41. Kent RJ, Crabtree MB, Miller BR. Transmission of West Nile Virus by Culex quinquefasciatus Say Infected with Culex Flavivirus Izabal. Tesh RB, editor. PLoS Negl Trop Dis. 2010;4: e671. doi:10.1371/journal.pntd.0000671.t003

42. Talavera S, Birnberg L, Nuñez AI, Muñoz-Muñoz F, Vázquez A, Busquets N. Culex flavivirus infection in a Culex pipiens mosquito colony and its effects on vector competence for Rift Valley fever phlebovirus. Parasit Vectors. 4 ed. BioMed Central; 2018;11: 405. doi:10.1186/s13071-018-2887-4

43. Brackney DE, Scott JC, Sagawa F, Woodward JE, Miller NA, Schilkey FD, et al. C6/36 Aedes albopictus Cells Have a Dysfunctional Antiviral RNA Interference Response. O’Neill SL, editor. PLoS Negl Trop Dis. 2010;4: e856. doi:10.1371/journal.pntd.0000856.t001

44. Lambrechts L, Quillery E, Noel V, Richardson JH, Jarman RG, Scott TW, et al. Specificity of resistance to dengue virus isolates is associated with genotypes of the mosquito antiviral gene Dicer-2. Proc R Soc B. 2012;280: 20122437–20122437. doi:10.1101/gad.1482006

45. Xi Z, Ramirez JL, Dimopoulos G. The Aedes aegypti toll pathway controls dengue virus infection. Schneider DS, editor. PLoS Pathog. 2008;4: e1000098. doi:10.1371/journal.ppat.1000098

46. Souza-Neto JA, Sim S, Dimopoulos G. An evolutionary conserved function of the JAK-STAT pathway in anti-dengue defense. Proc Natl Acad Sci USA. National Academy of Sciences; 2009;106: 17841–17846. doi:10.1073/pnas.0905006106

47. Sim S, Jupatanakul N, Ramirez JL, Kang S, Romero-Vivas CM, Mohammed H, et al. Transcriptomic Profiling of Diverse Aedes aegypti Strains Reveals Increased Basal-level Immune Activation in Dengue Virus-refractory Populations and Identifies Novel Virus-vector Molecular Interactions. Ribeiro JMC, editor. PLoS Negl Trop Dis. 2013;7: e2295. doi:10.1371/journal.pntd.0002295

48. Jupatanakul N, Sim S, Angleró-Rodríguez YI, Souza-Neto J, Das S, Poti KE, et al. Engineered Aedes aegypti JAK/STAT Pathway-Mediated Immunity to Dengue Virus. Olson KE, editor. PLoS Negl Trop Dis. Public Library of Science; 2017;11: e0005187. doi:10.1371/journal.pntd.0005187

49. Barletta A, Silva MCLN, Sorgine M. Validation of Aedes aegypti Aag-2 cells as a model for insect immune studies. Parasit Vectors. 2012;5: 148. doi:10.1146/annurev-biochem-060208-104626

50. Zhang R, Zhu Y, Pang X, Xiao X, Zhang R, Cheng G. Regulation of Antimicrobial Peptides in Aedes aegypti Aag2 Cells. Front Cell Infect Microbiol. Frontiers; 2017;7: 22. doi:10.3389/fcimb.2017.00022

51. Merkling SH, van Rij RP. Beyond RNAi: Antiviral defense strategies in Drosophila and mosquito. J Insect Physiol. Elsevier Ltd; 2013;59: 159–170. doi:10.1016/j.jinsphys.2012.07.004

52. Lan Q, Fallon M. Small heat shock proteins distinguish between two mosquito species and confirm identity of their cell lines. Am J Trop Med Hyg. 1990;43: 669–676.

53. Walker T, Jeffries CL, Mansfield KL, Johnson N. Mosquito cell lines: history, isolation, availability and application to assess the threat of arboviral transmission in the United Kingdom. Parasit Vectors. 2014;7: 382. doi:10.1186/1756-3305-7-382

54. Morazzani EM, Wiley MR, Murreddu MG, Adelman ZN, Myles KM. Production of virus-derived ping-pong-dependent piRNA-like small RNAs in the mosquito soma. Ding S-W, editor. PLoS Pathog. Public Library of Science; 2012;8: e1002470. doi:10.1371/journal.ppat.1002470

55. Stollar V, Thomas VL. An agent in the Aedes aegypti cell line (Peleg) which causes fusion of Aedes albopictus cells. Virology. 1975;64: 367–377.

56. Weger-Lucarelli J, Rückert C, Grubaugh ND, Misencik MJ, Armstrong PM, Stenglein MD, et al. Adventitious viruses persistently infect three commonly used mosquito cell lines. Virology. 2018;521: 175–180. doi:10.1016/j.virol.2018.06.007

57. Maringer K, Yousuf A, Heesom KJ, Fan J, Lee D, Fernandez-Sesma A, et al. Proteomics informed by transcriptomics for characterising active transposable elements and genome annotation in Aedes aegypti. BMC Genomics. BMC Genomics; 2017;18: 1–18. doi:10.1186/s12864-016-3432-5

58. Franzke K, Leggewie M, Sreenu VB, Jansen S, Heitmann A, Welch SR, et al. Detection, infection dynamics and small RNA response against Culex Y virus in mosquito-derived cells. J Gen Virol. 2018;99: 1739–1745. doi:10.1099/jgv.0.001173

59. Varjak M, Maringer K, Watson M, Sreenu VB, Fredericks AC, Pondeville E, et al. Aedes aegypti Piwi4 Is a Noncanonical PIWI Protein Involved in Antiviral Responses. Duprex WP, editor. American Society for Microbiology Journals; 2017;2: e00144–17. doi:10.1128/mSphere.00144-17

60. Varjak M, Donald CL, Mottram TJ, Sreenu VB, Merits A, Maringer K, et al. Characterization of the Zika virus induced small RNA response in Aedes aegypti cells. Olson KE, editor. PLoS Negl Trop Dis. Public Library of Science; 2017;11: e0006010. doi:10.1371/journal.pntd.0006010

61. Halstead SB, Simasthien P. Observations related to the pathogenesis of dengue hemorrhagic fever. II. Antigenic and biologic properties of dengue viruses and their association with disease response in the host. Yale Journal of Biology and Medicine. 1970;42: 276–292.

62. Dick GWA, Kitchen SF, Haddow AJ. Zika virus. I. Isolations and serological specificity. Transactions of the Royal Society of Tropical Medicine and Hygiene. 1952;46: 509–520.

63. Levine B, Goldman JE, Jiang HH, Griffin DE, Hardwick JM. Bc1-2 protects mice against fatal alphavirus encephalitis. Proc Natl Acad Sci USA. 1996;93: 4810–4815.

64. Hardwick M, Levine B. Sindbis virus vector system for functional analysis of apoptosis regulators. Methods in Enzymology. 2013;322: 1–17.

65. Stojdl DF, Lichty BD, tenOever BR, Paterson JM, Power AT, Knowles S, et al. VSV strains with defects in their ability to shutdown innate immunity are potent systemic anti-cancer agents. Cancer Cell. 2003;4: 263–275. doi:10.1016/S1535-6108(03)00241-1

66. Park MS, Shaw ML, Munoz-Jordan J, Cros JF, Nakaya T, Bouvier N, et al. Newcastle Disease Virus (NDV)-Based Assay Demonstrates Interferon-Antagonist Activity for the NDV V Protein and the Nipah Virus V, W, and C Proteins. J Virol. 2003;77: 1501–1511. doi:10.1128/JVI.77.2.1501-1511.2003

67. Scott JC, Brackney DE, Campbell CL, Bondu-Hawkins V, Hjelle B, Ebel GD, et al. Comparison of Dengue Virus Type 2-Specific Small RNAs from RNA Interference-Competent and –Incompetent Mosquito Cells. O’Neill SL, editor. PLoS Negl Trop Dis. 2010;4: e848. doi:10.1371/journal.pntd.0000848.t002

68. Anderson MAE, Gross TL, Myles KM, Adelman ZN. Validation of novel promoter sequences derived from two endogenous ubiquitin genes in transgenic Aedes aegypti. Insect Molecular Biology. 2010;19: 441–449. doi:10.1111/j.1365-2583.2010.01005.x

69. Schindelin J, Arganda-Carreras I, Frise E, Kaynig V, Longair M, Pietzsch T, et al. Fiji: an open-source platform for biological-image analysis. Nature Methods. Nature Publishing Group; 2012;9: 676–682. doi:10.1038/nmeth.2019

70. Whitfield ZJ, Dolan PT, Kunitomi M, Tassetto M, Seetin MG, Oh S, et al. The Diversity, Structure, and Function of Heritable Adaptive Immunity Sequences in the Aedes aegypti Genome. Curr Biol. 2017;27: 3511–3519.e7. doi:10.1016/j.cub.2017.09.067

71. Giraldo-Calderón GI, Emrich SJ, Maccallum RM, Maslen G, Dialynas E, Topalis P, et al. VectorBase: an updated bioinformatics resource for invertebrate vectors and other organisms related with human diseases. Nucleic Acids Research. Oxford University Press; 2015;43: D707–13. doi:10.1093/nar/gku1117

72. Goic B, Vodovar N, Mondotte JA, Monot C, Frangeul L, Blanc H, et al. RNA-mediated interference and reverse transcription control the persistence of RNA viruses in the insect model Drosophila. Nature Immunology. 2013;14: 396–403. doi:10.1038/ni.2542

73. Goic B, Stapleford KA, Frangeul L, Doucet AJ, Gausson V, Blanc H, et al. Virus-derived DNA drives mosquito vector tolerance to arboviral infection. Nat Commun. 2016;7: 12410. doi:10.1038/ncomms12410

74. Horst Ter AM, Nigg JC, Dekker FM, Falk BW. Endogenous viral elements are widespread in arthropod genomes and commonly give rise to piRNAs. J Virol. American Society for Microbiology Journals; 2018;: JVI.02124–18. doi:10.1128/JVI.02124-18

75. Palatini U, Miesen P, Carballar-Lejarazú R, Ometto L, Rizzo E, Tu Z, et al. Comparative genomics shows that viral integrations are abundant and express piRNAs in the arboviral vectors Aedes aegypti and Aedes albopictus. BMC Genomics. BioMed Central; 2017;18: 512. doi:10.1186/s12864-017-3903-3

76. Suzuki Y, Frangeul L, Dickson LB, Blanc H, Verdier Y, Vinh J, et al. Uncovering the repertoire of endogenous flaviviral elements in Aedes mosquito genomes. J Virol. American Society for Microbiology; 2017;: JVI.00571–17. doi:10.1128/JVI.00571-17

77. Crochu S, Cook S, Attoui H, Charrel RN, De Chesse R, Belhouchet M, et al. Sequences of flavivirus-related RNA viruses persist in DNA form integrated in the genome of Aedes spp. mosquitoes. J Gen Virol. 2004;85: 1971–1980. doi:10.1099/vir.0.79850-0

78. Mullane K, Williams M. Enhancing reproducibility: Failures from Reproducibility Initiatives underline core challenges. Biochem Pharmacol. 2017;138: 7–18. doi:10.1016/j.bcp.2017.04.008

79. van Cleef KWR, van Mierlo JT, Miesen P, Overheul GJ, Fros JJ, Schuster S, et al. Mosquito and Drosophila entomobirnaviruses suppress dsRNA-and siRNA-induced RNAi. Nucleic Acids Research. 2014;42: 8732–8744. doi:10.1093/nar/gku528

